# Natural and designed proteins inspired by extremotolerant organisms can form condensates and attenuate apoptosis in human cells

**DOI:** 10.1101/2021.10.01.462432

**Authors:** Mike T. Veling, Dan T. Nguyen, Nicole N. Thadani, Michela E. Oster, Nathan J. Rollins, Kelly P. Brock, Neville P. Bethel, Samuel Lim, David Baker, Jeffrey C. Way, Debora S. Marks, Roger L. Chang, Pamela A. Silver

**Author notes:** Co-first authors (Mike T. Veling, Dan T. Nguyen).

## Abstract

Many organisms can survive extreme conditions and successfully recover to normal life. This extremotolerant behavior has been attributed in part to repetitive, amphipathic, and intrinsically disordered proteins that are upregulated in the protected state. Here, we assemble a library of approximately 300 naturally-occurring and designed extremotolerance-associated proteins to assess their ability to protect human cells from chemically-induced apoptosis. We show that several proteins from tardigrades, nematodes, and the Chinese giant salamander are apoptosis protective. Notably, we identify a region of the human ApoE protein with similarity to extremotolerance-associated proteins that also protects against apoptosis. This region mirrors the phase separation behavior seen with such proteins, like the tardigrade protein CAHS2. Moreover, we identify a synthetic protein, DHR81, that shares this combination of elevated phase separation propensity and apoptosis protection. Finally, we demonstrate that driving protective proteins into the condensate state increases apoptosis protection, and highlight the ability for DHR81 condensates to sequester caspase-7. Taken together, this work draws a link between extremotolerance-associated proteins, condensate formation, and designing human cellular protection.

## INTRODUCTION

When exposed to extreme environmental stress, extremotolerance-associated (ExTol) organisms enter a state of reduced cellular metabolic activity that promotes survival in harsh conditions. This state has been shown to protect these organisms from high salinity, desiccation, and even the vacuum of outer space ^1^. To acclimate to these conditions, ExTol organisms significantly upregulate the production of protective metabolites, like trehalose ^2^, as well as tolerance-conferring proteins ^3^, many of which are intrinsically disordered proteins (IDPs) with repetitive sequence and predicted amphipathic helices.

While the mechanism of protection is unknown, one model suggests that at least some of these proteins mediate formation of a gel- or glass-like material state within an entire cell to protect it from damage ^3^. The model posits that such proteins promiscuously interact with membranes and other proteins, increasing intracellular viscosity and protecting vulnerable structures ^4-6^ (Figure 1A top). Such interactions would also work to prevent stress-induced protein unfolding and protein aggregation ^7^. This vitrification hypothesis is the subject of active investigation at the time of this writing, with evidence both for and against its validity ^8-10^. An alternative model suggests that these proteins form intracellular phase-separated membraneless compartments with distinct physical parameters (referred to here as condensates) that partition key stress-sensitive cellular components ^11^ (Figure 1A bottom). Within these dense protein structures, essential components are sequestered and protected from damage. The models are not mutually exclusive; for example, in the case of desiccation tolerance, continued dehydration could lead phase-separated compartments to fuse, comprise much (if not all) of the cell, and promote a glassy material state.

**Figure 1:**
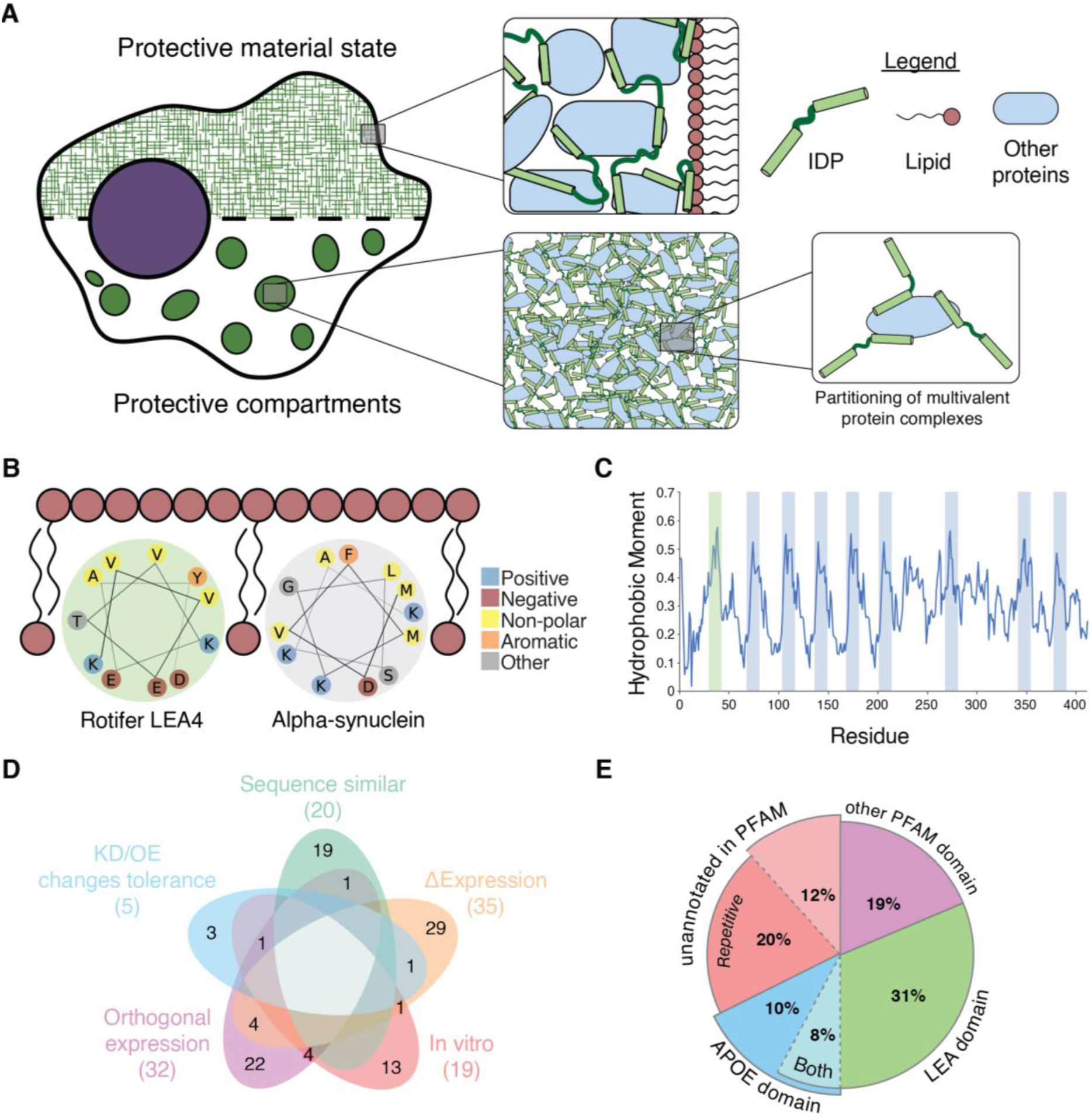
Selection of candidate extremotolerance-associated proteins (A) Illustration of different protective models for cellular stress tolerance. (B) Illustration of amphipathic membrane interaction for rotifer LEA4 protein and Alpha-synuclein, thus highlighting a feature indicative of extremotolerance-associated proteins. (C) Hydrophobic moment analysis of rotifer LEA4, showing the repetitive nature of the amphipathic signature. Highlighted green segment corresponds to the LEA4 helix in (B). (D) Venn diagram of the lines of evidence used to select targets in the final library. Abbreviations: KD, knockdown; OE, overexpression; Δ, change in expression levels. (E) Domain analysis of included targets reveals LEA and ApoE domains in almost half of the targets tested. The other half contains proteins with other PFAM domains (19%) or without any affiliated PFAM entry (32%). Of the proteins lacking a known PFAM annotation, many exhibit repetitive sequence (20% of all targets).

The resurgence of interest in biological phase separation, extensively reviewed in ^12-15^, has shown that human cells can employ similar strategies to survive stressful conditions. While the reversible formation of stress granules in healthy cells is the archetype of stress tolerance, recent studies have revealed phase separation as a key adaptation that cancer cells use to avoid apoptosis and cell death ^16-17^. Phase-separated condensates have been implicated in sequestering key tumor repressors^17^, enhancing or repressing gene expression in response to stress ^18^, and regulating mRNA or protein function through sequestration ^19^. These processes parallel the mechanisms that ExTol proteins are thought to use to protect complex biological systems, suggesting there could be some functional overlap. That is, sequence features and properties within ExTol proteins could be implicated in mammalian stress tolerance pathways, including apoptosis resistance.

A key sequence feature of some ExTol proteins is the presence of amphipathic helices ^20^. While disordered in most contexts, ExTol proteins can transiently form amphipathic helices in certain molecular environments, notably when interacting with and stabilizing membranes ^6^. The helices often exhibit a hydrophobic face, which interacts with core lipids within the membrane, flanked by positively charged residues that might interact with the negatively charged lipid head groups (Figure 1B). This amphipathic moment often repeats across the length of the protein (Figure 1C). This feature is a hallmark of the late embryonic abundant (LEA) class of ExTol proteins ^21^. Human membrane interacting proteins, like alpha-synuclein, also exhibit a similar repeat motif ^22^. Such parallels in sequence features suggest that orthogonal expression of ExTol proteins may confer their protective properties to other biological systems, like human cells.

Indeed, previous reports have shown that some of these ExTol proteins provide similar protective properties when expressed heterologously. Expressing tardigrade-specific IDPs (TDPs) in bacteria provides protection against desiccation ^3^. Anhydrin, an IDP upregulated by Aphelenchus avenae during dehydration, limits PABPN1-A17 aggregation, a hallmark of oculopharyngeal muscular dystrophy, when expressed in a human cell line ^7^. Similar reports demonstrate protection of human cells provided by expression of LEA proteins originating from the African midge ^23^ and by TDPs from tardigrades ^24^. These findings reinforce the promise of transferring protective phenotypes from their native contexts to human cells.

Taken together, these data indicate a potential avenue for using this class of proteins to protect biological systems in a controlled manner. To test this hypothesis, we generated approximately 300 sequences inspired by ExTol proteins and examined their ability to protect against apoptotic cell death. Our results show that human proteins containing sequence hallmarks of ExTol proteins as well as de novo designed condensate forming proteins can protect human cells from chemically-induced apoptosis. We also demonstrate that condensate induction using synthetic modifications can strengthen apoptotic protection. Finally, we show that key components of the apoptotic pathway can localize to inducible condensates, suggesting enzyme sequestration could be a mechanism of action.

## RESULTS

### Assembling a library of stress tolerance proteins

Based on the hallmark sequence features of ExTol proteins, we searched the published literature to identify proteins that not only contain these features but also exhibit some evidence for protection of biological structures. The library was generated based on over one hundred naturally-occurring proteins based on five lines of evidence for stress tolerance association, including: change in expression during extremotolerance; knockdown or overexpression changing the tolerant state; in vitro protection; orthogonal expression transferring tolerance; and sequence similarity to other proteins with these features (Figure 1D and Supplemental Table S1). We also examined our sequences for the presence of repetitive elements and intrinsic disorder (Supplemental Figure 1).

A PFAM domain analysis of the identified proteins reveals almost half of the identified targets have two key domains: Late Embryonic Abundant (LEA) and Apolipoprotein E (ApoE) domains (Figure 1E, Supplemental Table S1). LEA proteins are well known to be upregulated during the last stages of plant seed maturation ^25^. This domain is thought to protect the seed embryo during transport to its new destination. There is also evidence that this domain has been co-opted by bacteria as protection from desiccation and radiation ^26^. ApoE, in contrast, is not commonly discussed in the context of extremotolerance. ApoE proteins interact with lipid particles to allow for lipid reuptake into cells. Mutations within human ApoE have been identified as strong genetic predictors of Alzheimer’s disease in human populations ^27^. While seemingly unrelated to other proteins in our study, we chose to include human ApoE and its various mutations as ExTol proteins due to the prominence of the ApoE domain in the PFAM analysis and since they exhibit similar features as other selected proteins (e.g. their lipid interaction properties and phase separation tendencies ^28^).

### Designing novel extremotolerance-associated proteins

To test hypotheses about sequence features and their relationship to phase separation and apoptosis, we designed novel proteins based on the set of ExTol proteins. These new designs take 3 approaches (Figure 2): truncation of proteins to focus on repetitive, intrinsically disordered, and amphipathic regions; isolation of individual helices with desired sequence features; and de novo design of alpha helix-containing proteins that self-assemble into higher order structures. Through these designs, we sought to identify the minimal elements required for protection and phase separation.

**Figure 2:**
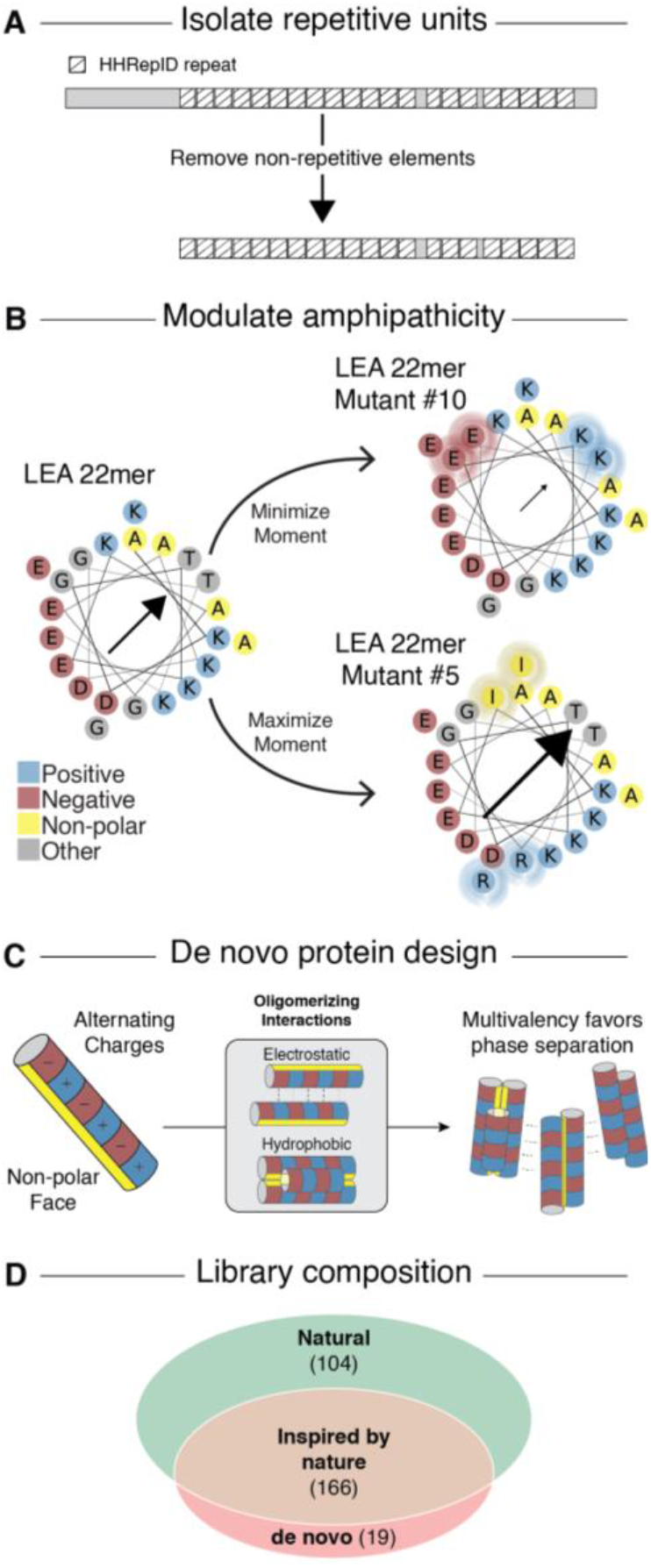
Design based on candidate extremotolerance-associated proteins (A) Truncations were made to isolate regions of repetitive and amphipathic sequence. (B) Small helices were selected and altered based on their hydrophobic moment. (C) Synthetic alpha helix-containing proteins designed to oligomerize. (D) Venn diagram highlighting the library that was tested.

First, we identified regions within natural proteins that contain intrinsic disorder ^29^, repetitive sequences ^30^, and amphipathic helices ^31^. Figure 2A illustrates the use of HHRepID on the sequence of full length human ApoE. Based on this analysis, we identified the repeating domain within ApoE for further testing. Other ExTol proteins were similarly modified and examined.

We also included short peptides based on the consensus sequence of repetitive regions that have previously been shown to be protective ^32^. To test if any of the sequence features we identified are important for apoptosis protection, we altered these features within these short peptides. For example, we altered the amphipathic character of these helices (Figure 2B) to determine if apoptosis protection was changed.

To examine whether generic condensate formation is sufficient to confer apoptosis protection, we also examined completely synthetic alpha helix-containing proteins that were designed to form repetitive structure with sequences orthogonal to naturally-occurring proteins ^33^. These designed helical repeat (DHR) proteins interact along the axis of the helix, forming long spiraling fibrils, but may also potentially self-interact along the face of the protein based on a pattern of positive and negative charges (Figure 2C). This ability to oligomerize and form multivalent complexes, which may then interact via weak non-covalent interactions, is an established determinant of condensate formation ^15^.

Together, our library consists of 104 naturally-occurring proteins, 166 truncations or isolations of helical regions, and 19 de novo designs (Figure 2D). These proteins interrogate several hypotheses about how disordered proteins may protect human cells against apoptosis.

### Apoptosis assay identifies extremotolerance-associated proteins that protect human cells

To model stress in a mammalian cell system, we induced cellular apoptosis by treatment with the chemotherapeutic camptothecin (CPT), a DNA topoisomerase inhibitor well known to trigger apoptosis ^34-35^. Under these conditions, protection against apoptosis could be achieved through a number of avenues, including protection against the DNA damage caused by CPT-induced topoisomerase inhibition ^34^, interruption of the signaling cascade, or even protection of the mitochondrial membrane whose rupture ultimately triggers apoptosis ^36^.

To test our designs for protection against apoptosis, we used a human cell-based apoptosis assay (Figure 3A). This assay uses HT-1080 fibrosarcoma cells from ATCC ^37^. These cells are seeded in a 96-well plate and allowed to adhere overnight. The next day, the cells are co-transfected with a plasmid encoding the ExTol candidate proteins of interest as well as a transfection marker. The following day, the media is swapped to one containing 13.2 μM CPT. After addition of CPT, caspase-3 and −7 activation is measured by microscopy every 45 minutes for 1 day using a fluorophore quencher pair attached to either end of a peptide with the sequence “DEVD”. As caspase-3 and −7 are activated, the DEVD peptide is cleaved, releasing the fluorophore from the quencher. The unquenched fluorophore localizes to the nucleus where it can be quantified by imaging on a single-cell level. The single-cell resolution allows for the separation of transfected and non-transfected cells based on the transfection marker. Only single cells with the transfection marker were subjected to further analysis; Figure 3B shows sample images highlighting the type of data being analyzed.

**Figure 3:**
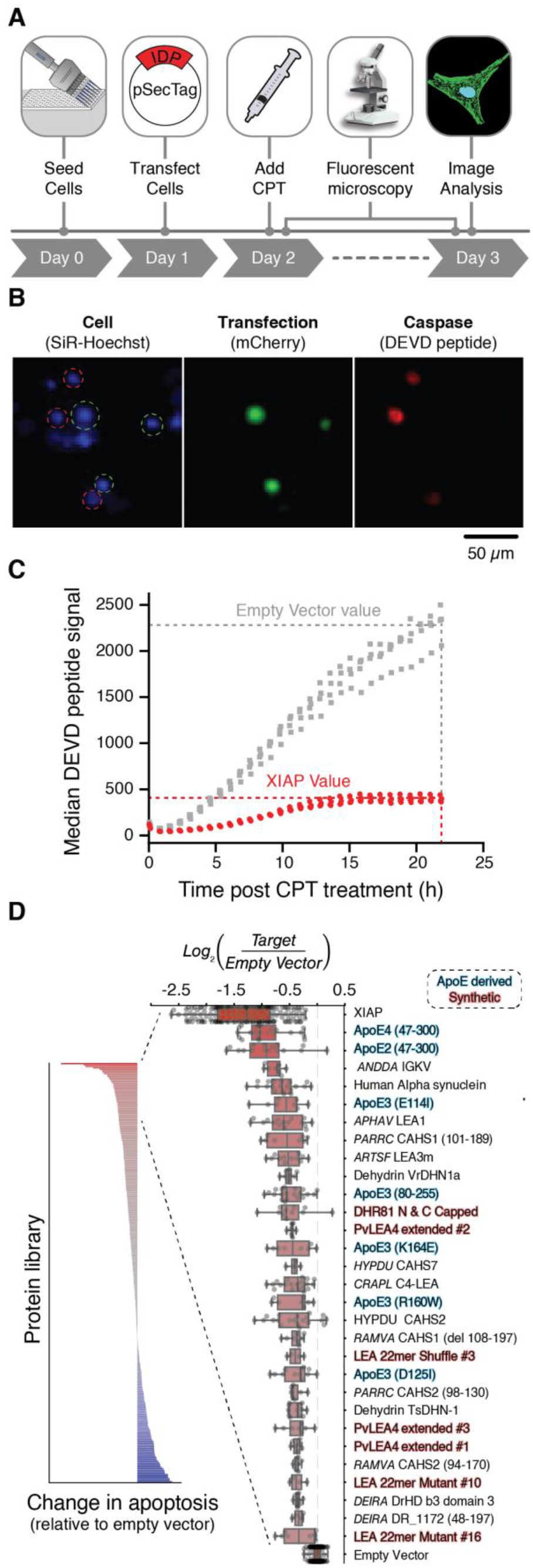
Screen for CPT resistance by extremotolerance-associated proteins (A) Flow diagram of the assay. (B) Representative images showing cells during the assay. Cells are identified using SiR-Hoechst marking the nucleus (blue). Transfected cells are identified by the presence of the mCherry marker (green). Finally, the degree of apoptosis is identified by the presence of the cleaved DEVD peptide (red). (C) Positive control showing apoptosis reduction upon expression of XIAP. (D) Bar graph highlighting the apoptosis resistance provided by different constructs, including zoomed in area to highlight the top hits in the assay. Source species of a sequence are abbreviated and shown in italics; abbreviations are drawn from Uniprot, including RAMVA (Ramazzottius varieornatus), HYPDU (Hypsibius dujardini or Hypsibius exemplaris historical naming), PARRC (Paramacrobiotus richtersi), ANDDA (Andrias davidianus), APHAV (Aphelenchus avenae), ARTSF (Artemia franciscana), CRAPL (Craterostigma plantagineum), and DEIRA (Deinococcus radiodurans).

On every plate, an empty vector negative control is run as well as a positive control containing the X-linked inhibitor of apoptosis (XIAP). This protein is known to bind to caspase-3 and −7 to block their activity ^38^. Figure 3C shows example data comparing median DEVD peptide signal intensity over time after induction with CPT of four wells on the same plate transfected with XIAP and empty vector. The results show a reduction in DEVD peptide signal intensity when the XIAP-containing vector is transfected, demonstrating that the assay can identify modulators of apoptosis.

To convert these time-resolved data to a measure of protection against apoptosis, the maximum DEVD peptide signal intensity is calculated and normalized to a plate-matched empty vector control (see methods for further details). The normalized apoptosis induction intensity is shown in Figure 3D left. The data are slightly skewed towards apoptosis-protection, which supports our hypothesis that the targeted proteins can protect against apoptosis.

To highlight the top-performing designs, the top 10% of the targets are shown in Figure 3D right. As expected, the XIAP positive control showed the strongest anti-apoptotic effect. The two top-performing hits were ApoE designs. Human alpha-synuclein is the 5^th^ hit, which is expected as it is known to be both intrinsically disordered and anti-apoptotic ^39^. Additionally, several ApoE designs and short LEA helices are top hits. The data used to generate these figures is included in Supplemental Table S2.

### Known extremotolerance-associated proteins affect apoptosis

Among the top hits in apoptosis reduction are an IGKV protein highly expressed in mucus of the Chinese giant salamander Andrias davidianus ^40^ that is used as a medical glue in traditional Chinese medicine, an LEA protein from the nematode Aphelenchus avenae previously shown to have the ability to reduce a form of protein aggregation associated with oculopharyngeal muscular dystrophy in human cells ^7^, and a mitochondrial LEA protein from the brine shrimp Artemia franciscana shown to protect human cells during desiccation ^23^ (Figure 3D). The prominence of these ExTol proteins suggests the assay can help identify other proteins that can protect cells from more general stresses beyond CPT-induced apoptosis.

### Short repetitive peptides and de novo designs modulate apoptosis

LEA repeat-containing peptides are among the top-performing hits in the screen. While the original short LEA designs from Furuki et al. 2019 had a minor effect reducing apoptosis, several of the designs tested here performed better (Supplemental Figure 2A) ^32^. As 22 amino acids is small and could lead to peptide stability issues, longer 50-amino acid designs inspired by the same LEA proteins utilized by Furuki et al. were also developed (Supplemental Figure 2B). Some of these designs were also able to reduce apoptosis, including the PvLEA4 with extended repeats (notably PvLEA4 repeats extended #2), which fell into the top hits of the entire library (Figure 3D and Supplemental Figure 2B). These results suggest that optimization of synthetic designs yields proteins with stronger function than naturally-occurring proteins.

While these proteins were designed to interrogate the relationship between sequence and apoptosis protection in short peptides, little relationship was found with respect to the sequence features we analyzed (Supplemental Figure 2C). Several designs were made that would alter the hydrophobic moments of putative helices these peptides are predicted to form. To examine if this sequence property relates to the stress-tolerance function of these proteins, we attempted to correlate this property with assay performance. However, no significant correlation was found, suggesting (at least in these cases and experimental conditions) that degree of amphipathicity on its own is not the key determinant of stress-tolerance function of these proteins.

### The repeat region of ApoE comprises its minimal anti-apoptotic unit

The top two hits in the assay were ApoE4 (47-300) and ApoE2 (47-300). These truncated forms of ApoE maintain the repeat domains shown in Figure 4A, while removing the signal peptide and terminal regions. This suggests that the repeat region of ApoE accounts for its core anti-apoptotic functionality. Interestingly, the full length ApoE2 and E4 do not show up as top hits, suggesting that these truncated forms improve on the apoptosis protection provided by the full-length variants (Figure 4B).

**Figure 4.**
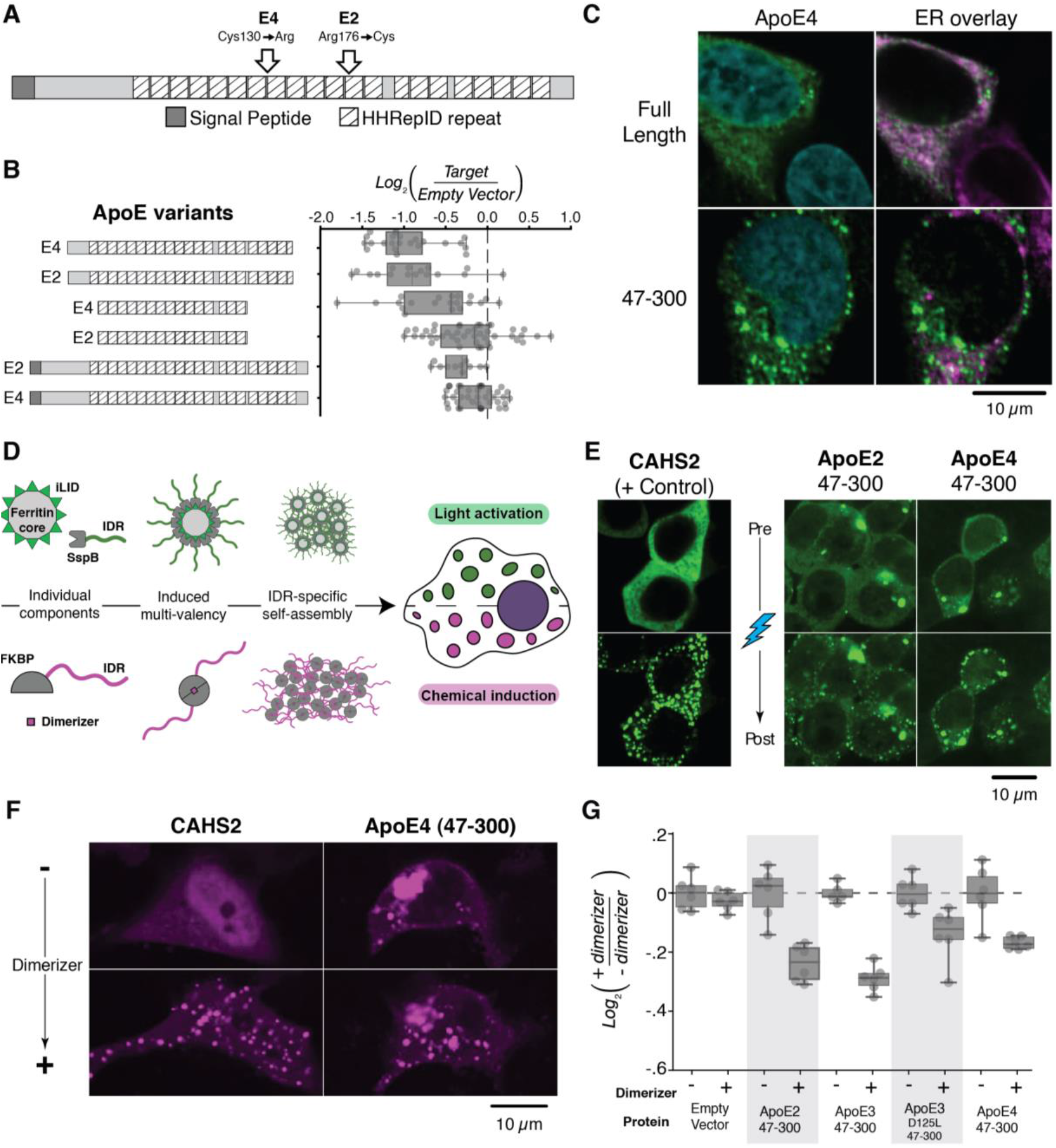
ApoE truncations and synthetic condensates further reduce apoptosis. (A) Gene diagram highlighting the repetitive region of ApoE. (B) Boxplots highlighting the apoptosis assay performance of the full length and truncated forms of ApoE2 and E4. (C) Immunofluorescence of ApoE4, full length and truncated forms. White indicates colocalization, which is more apparent with the full-length protein. The truncated form appears to exhibit larger and more distinguishable puncta. (D) Diagram of the induced multi-valency systems implemented here, including the light-inducible system (top) and the chemically-inducible system (bottom). For the chemically-induced system, we used two FKBP domains. (E) Condensate formation assay using the light-inducible system, including the PARRC CAHS2 positive control showing condensates only upon light-induction and several ApoE examples that form condensates both before and after induction. (F) Condensate formation assay using the chemically-inducible system, including the PARRC CAHS2 positive control showing condensates only upon added dimerizer and ApoE4 (47-300) exhibiting condensates both before and after induction. (G) Apoptosis data showing reduction in apoptosis upon condensate induction with dimerizer. Data are normalized to mean apoptosis level of uninduced (i.e. no dimerizer) sample for each construct.

Based on the location of the signal peptide, it is expected that ApoE4 should localize to the secretory pathway whereas the truncated form may be cytoplasmic. To test this hypothesis, cells were subjected to Western blot and immunofluorescence analysis using commercially available antibodies (Abcam ab24139; Figure 4C and Supplemental Figure 3A). The Western blot results show that both the full-length and truncated forms of ApoE4 are expressed at the expected size (Supplemental Figure 3A). The immunofluorescence shows co-localization between the ApoE4 full length protein and the endoplasmic reticulum (marked by Thermo’s PDI antibody MA3-019, Figure 4C). However, the truncated form did not show the same co-localization, suggesting as expected that it may not enter the secretory pathway as the full-length version does.

Strikingly, the truncated version appeared to form more pronounced puncta throughout the cytoplasm of the cell, suggesting the formation of potentially phase-separated condensates. In screening through the point mutants of ApoE also tested here, we found a similar condensate phenotype with the full length ApoE3 D125I mutation (Supplemental Figure 3B). Together, these data suggest that ApoE may be able to support condensate formation and confer apoptosis protection in part via the protective compartment mechanism (Figure 1A bottom).

### Induced multivalency to probe the relation between condensates and apoptosis

We made use of established condensate engineering tools ^41^ to test for latent phase separation propensity among a selection of ExTol proteins representing the range of apoptosis effects. We used both light- and chemically-inducible systems of multivalency. In the light system, a ferritin core (comprised of self-assembled monomers, FTH1 domain-eGFP-iLID) becomes decorated with SspB-modified proteins upon exposure to blue light ^42^ (Figure 4D top). In the chemical system, multivalency (i.e. multiple arms of the ExTol protein of interest) is induced using a dimerization agent ^43^ that brings together FKBP-modified proteins ^41^ (Figure 4D bottom). These inducible methods increase the valency of a system, and thus work to favor phase separation. In this way, they can help reveal any latent ability to form condensates that is not immediately observable without modification.

Condensates were indeed observed in a number of ExTol proteins (Supplemental Figure 4), including ApoE, ARTSF LEA3m, DHR81, HYPDU CAHS3, and PARRC CAHS2 using the light-inducible system. For those proteins with no visible condensates, the experimental conditions may be insufficient to support phase separation. PARRC CAHS2 exhibited the most dramatic phenotype, which is consistent with recent work describing the gel forming capabilities of CAHS proteins ^44-45^ (Figure 4E&F). ApoE variants were of interest, based on the puncta observed in the top performing ApoE constructs. All versions of ApoE tested formed puncta without activation, consistent with our previous observation using untagged native ApoE (Figure 4C). Upon light- or chemical-activation, there is a noticeable increase in condensates, exemplifying how phase separation can be enhanced with increased multivalency (Figure 4E&F).

To examine how enhanced phase separation may affect apoptosis, we performed the apoptosis assay with ExTol proteins that were modified for chemically-induced multivalency. Chemical induction was more compatible with the apoptosis assay, while enabling the same increase in condensates seen with the light-induced system (Figure 4F). We observed that the presence of the chemical dimerizer had little to no effect on apoptosis in cells transfected with the empty vector, with a construct encoding the dimerization domains alone, or with constructs encoding an ExTol protein that was not strongly anti-apoptotic (e.g. PARRC CAHS2) (Figure 4G; Supplemental Figure 5). In contrast, with constructs encoding a strongly anti-apoptotic ExTol protein – including ApoE, ARTSF LEA3m, and DHR81 – added dimerizer corresponded with reduced apoptosis in transfected cells (Figure 4G; Supplemental Figure 5). These data suggest that condensate formation plays a role in the anti-apoptotic effect of these proteins.

### Synthetic DHR81 condensates sequester caspase-7

To begin elucidating the mechanism by which ExTol proteins protect against apoptosis, particularly the hypothesis of membraneless compartments that sequester essential effectors, we examined the ability for the various condensates to partition apoptotic signaling molecules, identifying the synthetic protein DHR81 as the only ExTol protein capable of doing so. DHR81 is a synthetic alpha helical-rich protein designed to promote self-interaction. It is formed from four helical hairpin repeat regions; the core of the protein is hydrophobic to ensure consistent folding, while its outside is decorated with checkerboard positive and negative charges (Figure 5A) ^33^. This structure ensures the protein is well folded, while also enabling the potential for higher order structure formation via electrostatic interactions. Not only did DHR81 support condensates in the chemically-inducible condensate system (Figure 5B), it also exhibited a strong increase in apoptotic protection after induced-multivalency (Figure 5C; Supplemental Figure 5&6A).

**Figure 5.**
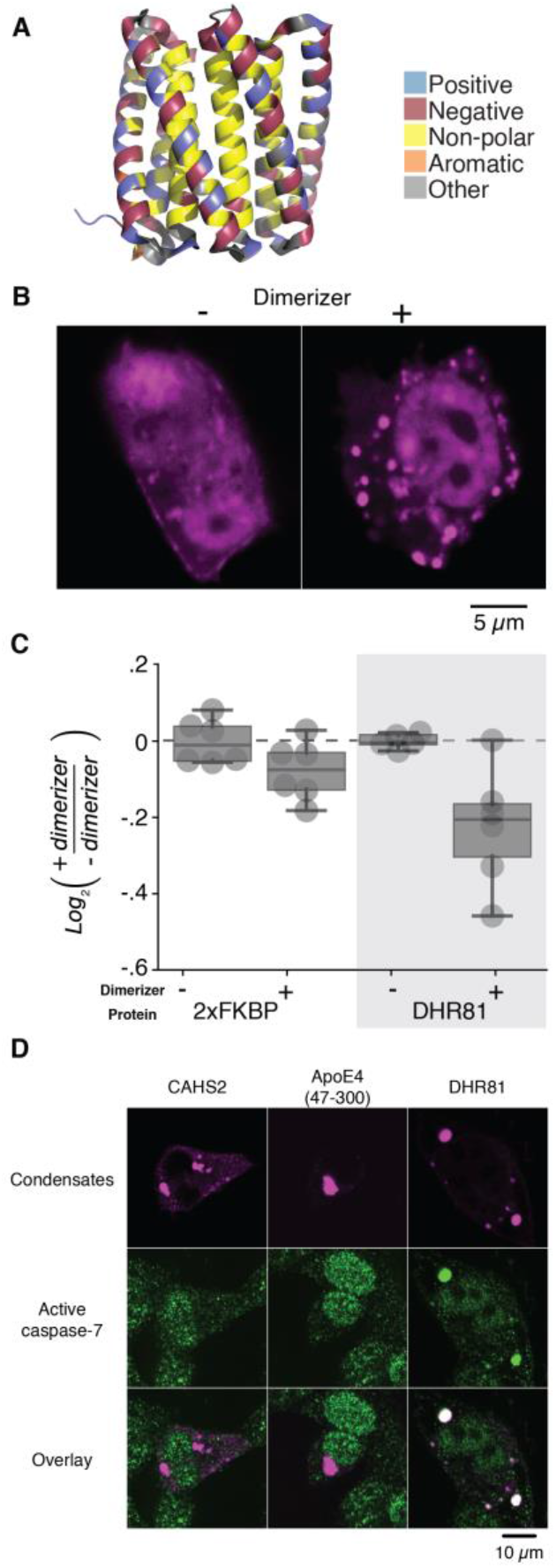
DHR81 condensate formation and apoptosis reduction. (A) Diagram of DHR81, illustrating a charge distribution that can support oligomerization. (B) Sample images highlighting how the synthetic protein DHR81 supports chemically-induced condensates. (C) Apoptosis is further reduced in the presence of dimerizer (i.e. with induced DHR81 condensates). The FKBP-modifications alone, which enable chemically-induced multi-valency, have a minimal effect on apoptosis. (D) Activated caspase-7 (green) co-localizes with DHR81 condensates but not PARRC CAHS2 or ApoE4 (47-300) condensates (magenta). White indicates colocalization.

As such, we chose DHR81, along with a few other ExTol proteins with notable phase separation phenotypes (e.g. ApoE4 and PARRC CAHS2), to further explore how their condensates may confer an anti-apoptotic effect. Specifically, we used immunofluorescence to examine whether they sequester major components of apoptotic signaling, namely caspase-3 and −7. The results showed that activated caspase-7 localized to DHR81 condensates but not PARRC CAHS2 or ApoE4 condensates (Figure 5D; Supplemental Figure 5C), while caspase-3 did not localize to any of the examined condensates (Supplemental Figure 6B). Even though condensate properties may confound (i.e. physically occlude) antibody binding during immunofluorescence, the positive DHR81 result suggests sequestration of key apoptotic factors can indeed be a potential mechanism by which these condensates reduce apoptosis.

## Discussion

To broadly evaluate the ability of ExTol proteins to protect orthogonal systems from environmental stress, we studied over one hundred different ExTol proteins. We used these as a basis to design approximately 300 variants, including truncations and fusion proteins. These designs were evaluated on their ability to protect human HT-1080 cells against CPT-induced apoptosis. The results showed that known ExTol proteins, ApoE variants, and the synthetic DHR81 protein could modulate apoptosis levels when transferred to human cells.

Protection against apoptosis is a key feature of many ExTol proteins. As apoptosis is a common cellular response to stress, the ability to mitigate this response is important in providing cell-autonomous protection in stressful environments. While the mechanism resilient organisms use to survive is still not fully understood, it is becoming increasingly clear that phase separation may play an important role. Some ExTol proteins are proposed to form expansive higher order structures (e.g. gels or glassy systems) that reduce the viscosity of the cytoplasm, especially in desiccated states ^3^. Other reports show that these proteins could mediate formation of protective phase-separated compartments (i.e. condensates) that sequester damage-inducing proteins and/or protect important targets ^11^. Based on our data, it appears the latter may be at play for proteins like ApoE, ARTSF LEA3m, and DHR81, which exhibit anti-apoptotic effects while also being condensate prone.

Condensates alone are not sufficient for apoptosis protection. While we observe condensate formation in many of our top performing hits, we also observe condensate formation in targets like HYPDU CAHS3 and PARRC CAHS2, which do not reduce apoptosis very much. Additionally, some proteins like PARRC CAHS1 (101-189) are truncated in ways predicted to reduce condensate formation^44, 46-47^, but they still perform well in apoptosis protection. These observations suggest yet-to-be-discovered mechanisms are likely responsible for apoptosis protection in these cases. However, for those proteins that exhibit both anti-apoptotic effects as well as elevated phase separation propensities, increasing the amount of condensates via an engineered inducible system led to stronger apoptosis protection (Supplemental Figure 5). In these cases, condensates could be a key part of the protective mechanism.

Specifically, we observed that activated caspase-7 partitions into DHR81 condensates, suggesting caspase sequestration as a potential anti-apoptotic mechanism. Phase-separated condensates and higher order assemblies are already implicated in mediating apoptosis: stress granules inhibit apoptosis by suppressing MAPK pathways ^48^; the apoptosome relies on Apaf-1 multimerization, which has been reported to undergo amyloid fibrillation in specific conditions ^49^; and stress-dependent phase separation modulates proteasome formation, whose function affects apoptosis ^50-51^. It is interesting to note that we only observe clear caspase-7 partitioning into DHR81 condensates, suggesting the possibility of a specific binding interaction in this case. ApoE and ARTSF LEA3m condensates may be sequestering other apoptotic factors or using a different mechanism to reduce apoptosis.

It is also possible that these ExTol proteins achieve their anti-apoptotic effect by implementing a more general physical mechanism, namely slowing the intracellular mobility of biological effectors. There are various reports describing how the effective viscosity within the cytoplasm can be increased in response to external stresses ^4-5^ or mediated by protein composition, specifically intrinsically disordered proteins ^52^. Notably, a pervasive cell-spanning gel or network is not necessary to achieve such an effect; it is possible that the presence of condensates, coupled with their ability to selectively partition biomolecules (i.e. control what biomolecules can enter and diffuse through them), could broadly slow biomolecular movement. That is, by forcing different proteins to either be trapped in them or diffuse around them, condensates can decrease protein mobility and perturb e.g. signaling pathways. Our results showing a lack of caspase-3 and −7 sequestration by most of our protein condensates could thus support this more general mechanism of condensates perturbing biology, though they certainly do not disqualify sequestration of other effectors not examined in this study.

Condensates are becoming increasingly implicated in disease ^13^, particularly when their structural metastability and reversibility is dysregulated (e.g. through mutation). Notably, drugs that block the formation of amyloid fibrils and plaques, which are products of phase separation dysregulation, are promising yet strongly debated ^53^ therapies for many neurological diseases. As such, our observations concerning ApoE proteins, while unexpected from an extremotolerance perspective, are particularly salient. We found that expression of a cytoplasmic form of ApoE is protective against apoptosis. In particular, the truncated repeat region of ApoE is sufficient and shows the strongest anti-apoptotic activity of all designs we tested. Others have shown similar effects for exogenously added ApoE-containing liposomes protecting rat neurons against apoptosis ^54^, though we show the effect through overexpression of ApoE inside of human cells. When combined with our condensate results and the clinical relevance of ApoE in Alzheimer’s disease ^55^, these observations suggest the anti-apoptotic effect observed here warrants further study, to further elucidate its role in Alzheimer’s disease progression.

Altogether, our work expands upon the understanding of the relationship between condensate formation and cell physiology, demonstrating how condensate accumulation can be beneficial in reducing apoptosis while also identifying diverse new protein domains that confer protection. The proteins and concepts presented here demonstrate the utility of this class of proteins for engineering new properties into cells and ultimately higher organisms.

## Methods

### Cell culturing and maintenance

HT-1080 cells (ATCC, CCL-121) were maintained in EMEM medium supplemented with 10% FBS and Penicillin-Streptomycin according to ATCC’s protocol. For additional details see Supplemental Methods.

### DNA Preparation

Apoptosis constructs were synthesized by Twist Bioscience and transformed into DH5α cells for amplification and plasmid purification. Condensate assay constructs were subcloned from these synthesized constructs into vectors containing the appropriate chemically- or light-inducible condensate elements. DNA was prepared using the QIAGEN Plasmid Plus Midi Kit (Qiagen 12943) for transfection grade DNA purification. See Supplemental Methods for additional details.

### Apoptosis assay

200 μL of HT-1080 cells at 7.5×10^4^ cells per mL (15000 cells per well) were added to each well of a microscope grade 96 well plate (Corning 3603). Plates were then centrifuged at 330 rpm for 3 min (Eppendorf, Centrifuge 5810). Plates were then incubated at 37°C for 24 hours. Cells were then transfected with appropriate DNA constructs with the lipofectamine 3000 kit (Thermo Fisher Scientific, L3000015) as specified by the manufacturer. 24 hours after transfection, the transfection media was removed and replaced with FluoroBrite DMEM Media (Thermo A1896702) supplemented with 1x GlutaMAX (Thermo 35-050-061) and 10% FBS (Thermo Fisher Scientific, 10082147). This media also contained 13.2 μM (S)-(+)-Camptothecin (Sigma C9911-100MG), 0.3 μM SiR-DNA stain (SpiroChrome CY-SC007), and 1.25 μM of the DEVD peptide for detecting caspase activation (Sartorius 4440). The plates were sealed with sterile Breathe-Easy film (USA scientific 9123-6100) and incubated at 37°C with 5% CO_2_ for 30 minutes. After 30 min incubation, the plates were placed in an ImageXpress-confocal plate imager (Molecular Devices) at 37°C with 5% CO_2_ and saturated humidity through the presence of a heated water bath in the chamber. Images of all wells were acquired every 45 minutes and stored on a remote server for later analysis. See Supplemental Methods for further details on this method as well as the image analysis pipeline used.

### Confocal microscopy

Confocal microscopy was performed at the Nikon Imaging Center at Harvard Medical School using a Nikon Ti fluorescence microscope attached to a Yokogawa W1 spinning disk confocal. See Supplemental Methods for further details on the equipment and settings used.

### Light-inducible condensate assay

HEK293-T cells were seeded on 96 well glass bottom plates from Cellvis (P96-1.5H-N). Cells within each well were transfected with 50 ng of the LLPS1-iLID::EGFP::FTH1 (Addgene plasmid #122149) construct containing the core FTH1 scaffolding domain attached to iLID for light-inducible interactions. The cells were also transfected with 50 ng of a construct containing the target protein fused to mCherry and SspB. The transfection was allowed to continue overnight before the media was swapped to FluoroBrite media supplemented with 10% FBS and 1x GlutaMAX. Pre-induction images were acquired using the settings listed in the Supplemental Methods for mCherry and eGFP. The cells were then exposed to 1 second of the DAPI imaging protocol described in Supplemental Methods before new images were captured on the mCherry and eGFP settings.

### Chemically-inducible condensate assay

HT-1080 cells were seeded on 96 well glass bottom plates from Cellvis (P96-1.5H-N). Cells within each well were transfected with 50 ng of target construct fused to a mCherry-2xFKBP domain. The transfection was allowed to continue overnight before the media was swapped to FluoroBrite media supplemented with 10% FBS, 1x GlutaMAX, and 300 nM SiR-Hoechst. The cells were then imaged using the settings listed in the Supplemental Methods for mCherry and SiR-Hoechst for the pre-induction images. Then, B/B dimerizer (AP20187 ligand; TaKaRa 635058) was added to a final concentration of 500 nM in the well. Imaging continued for 15 minutes with new images every 45 seconds.

### Caspase assay with chemically-induced condensates

These caspase assays are performed similarly to the standard caspase assays with a few key differences. Briefly, HT-1080 cells were seeded as normal on to 96 well plates and allowed to adhere overnight. The next day, 12.5 ng of nuclear mCherry marker and 37.5 ng of the target construct fused to mCherry-2xFKBP was transfected into the cells as described above. The next day, 500 nM of the B/B dimerizer (TaKaRa 635058) was added 2 hours before the start of the assay (or an Ethanol vehicle control was added). The media was then removed and replaced with the apoptosis assay media described above with the addition of 500 nM B/B dimerizer or a vehicle control. See Supplemental Methods for further details.

## Abbreviations

ExTol: extremotolerance-associated
IDP: Intrinsically Disordered Protein
LEA: Late Embryonic Abundant (protein)
ApoE: Apolipoprotein E
DHR: Designed helical repeat (protein)
CPT: Camptothecin
ATCC: American Type Culture Collection
XIAP: X-linked inhibitor of apoptosis
CAHS: Cytosolic Abundant Heat Soluble (protein)
FTH1: Ferritin Heavy Chain 1
iLID: improved light inducible dimer
SspB: Stringent Starvation Protein B
eGFP: Enhanced Green Fluorescent Protein
FKBP: FK506 binding protein
PvLEA: Polypedilum vanderplanki Late Embryonic Abundant (Protein)
MAPK: Mitogen-activated protein kinase
Apaf: Apoptotic protease activating factor
EMEM: Eagle’s Minimum Essential Medium
FBS: Fetal bovine serum
DMEM: Dulbecco’s Modified Eagle Medium

## Acknowledgements

Research was sponsored by the NSF, DARPA, the Wyss Institute for Biologically inspired engineering, and the Army Research Office and was accomplished under Cooperative Agreement Number W911NF-19-2-0017. M.T.V. was sponsored by the National Science Foundation under Grant No. 2010370. D.T.N., N.N.T., M.E.O., N.J.R., K.P.B., J.C.W, D.S.M., and P.A.S. were supported by the Harvard Medical School Department of Systems Biology. J.C.W. was further supported by the Harvard Medical School Laboratory of Systems Pharmacology. K.P.B. and D.S.M. were further supported by CZI Neurodegenerative Challenge Ben Barres Early Career Award (CZI2018-191853). N.P.B was further supported by the HHMI Hanna Gray fellowship (grant GT11817). We would like to acknowledge Clarence Yapp for help with microscopy as well as the NCI U54-CA225088 grant and a gift from the Ludwig Institute for Cancer Research to the Laboratory for Systems Pharmacology. We would like to acknowledge the Nikon Imaging Center at Harvard Medical Center for providing microscope support. The views and conclusions contained in this document are those of the authors and should not be interpreted as representing the official policies, either expressed or implied, of any funding agency including the NSF, DARPA, the Army Research Office, or the U.S. Government. The U.S. Government is authorized to reproduce and distribute reprints for Government purposes notwithstanding any copyright notation herein. Visual table of contents entry was created with support from BioRender.com.

## Author Information

Correspondences should be sent to: Pamela Silver, Harvard Medical School Department of Systems Biology 200 Longwood Avenue Warren Alpert Building Boston, MA 02115; and to Roger Chang, Albert Einstein College of Medicine Department of Systems & Computational Biology 1301 Morris Park Avenue Price Center Bronx, NY 10461.

## Author Contributions

M.T.V. and D.T.N. designed and performed experiments, analyzed the data, and wrote the paper. N.N.T. analyzed the data and wrote the paper. M.E.O provided technical support. N.J.R., K.B.B., N.P.B., and S.L., designed constructs. D.B. and J.C.W. managed the teams and provided key insight. D.S.M., R.L.C., and P.A.S., designed constructs, managed teams, and wrote the paper.

## Supporting information

<u>Supplemental figures and table descriptions:</u> Contains Supplemental Figures 1 - 6 referenced in the main text, as well as full descriptions of the Supplemental Tables. This includes an analysis of the disorder and repetitive distribution of the identified ExTol proteins, extended analysis of the relationship between the synthetic LEA proteins and apoptosis protection, additional data on the ApoE Western blot and immunofluorescence experiments, data regarding our screen for condensate formation across the spectrum of apoptosis related constructs, a broader look at condensate formation as it relates to apoptosis protection and condensate sequestration, and extended data on DHR81.

<u>Supplemental Methods:</u> Contains detailed information about the methodology used in this study.

<u>Supplemental Table S1:</u> Reports sequences of all the proteins and designs discussed in this work.

<u>Supplemental Table S2:</u> Reports all normalized apoptosis data collected throughout this work.

## For Table of Contents Use Only

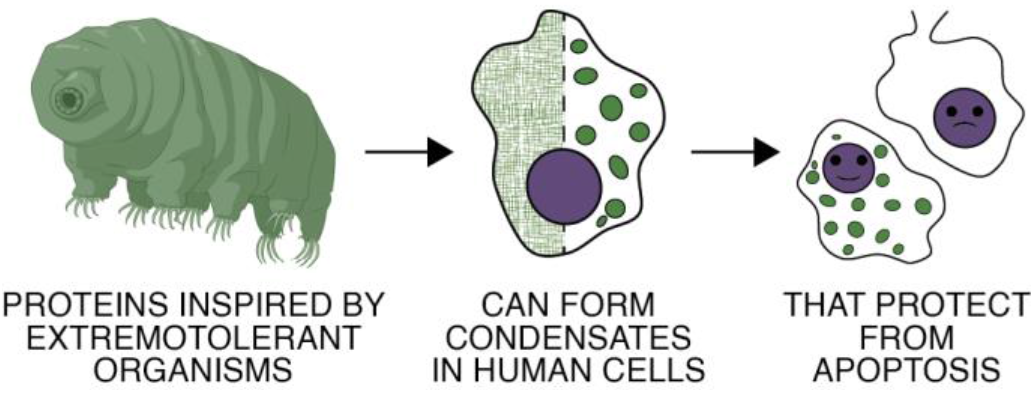

