## Supplemental Figures for "Natural and designed proteins inspired by extremotolerant organisms can form condensates and attenuate apoptosis in human cells"

1. Department of Systems Biology, Harvard Medical School, Boston, MA, 02115, USA;
2. Wyss Institute for Biologically Inspired Engineering, Boston, MA, 02115, USA;
3. Institute for Protein Design, University of Washington, Seattle, WA, 98195, USA;
4. Department of Biochemistry, University of Washington, Seattle, WA, 98195, USA;
5. Howard Hughes Medical Institute, University of Washington, Seattle, WA, 98195, USA;
6. Broad Institute of Harvard and MIT, Cambridge, MA, 02142, USA;
7. Department of Systems & Computational Biology, Albert Einstein College of Medicine, Bronx, NY, 10461, USA;
8. Co-first authors

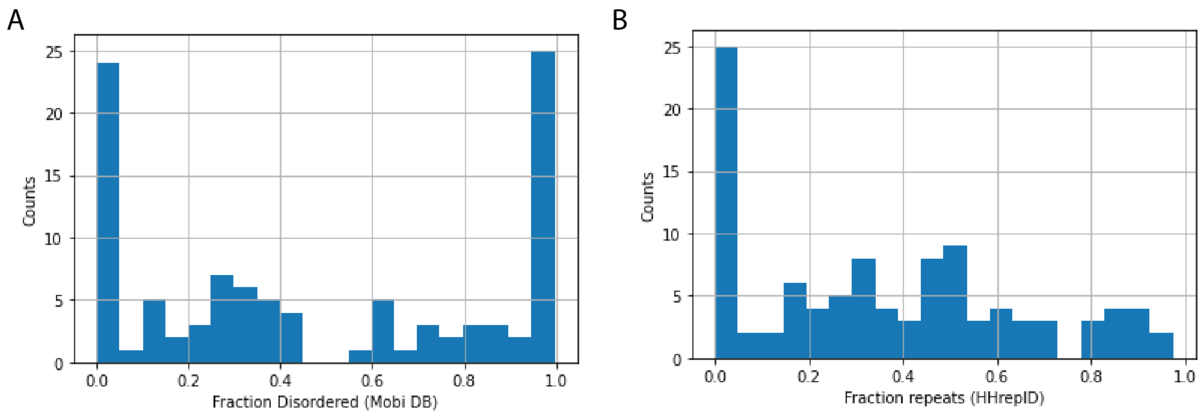


**Supplemental Figure 1** (A) Distribution of fractional disorder within the sequences of the selected ExTol proteins. (B) Distribution of fractional repeats within the sequences of the selected ExTol proteins.

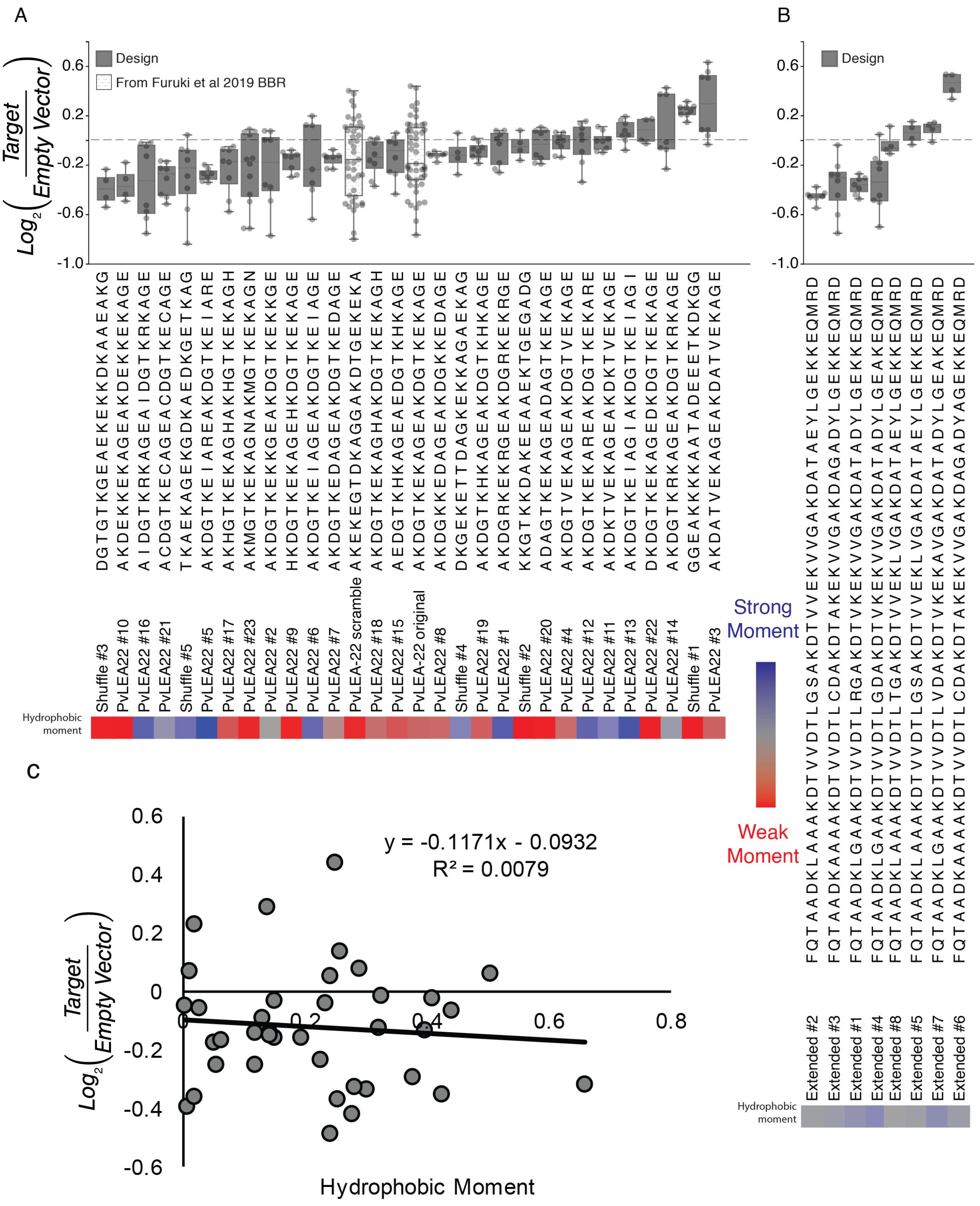


**Supplemental Figure 2** Graph of various LEA repeats tested (A) and extended LEA repeats tested (B), with full sequences and performance. (C) Correlation analysis for relationship between hydrophobic moment with apoptosis assay performance.

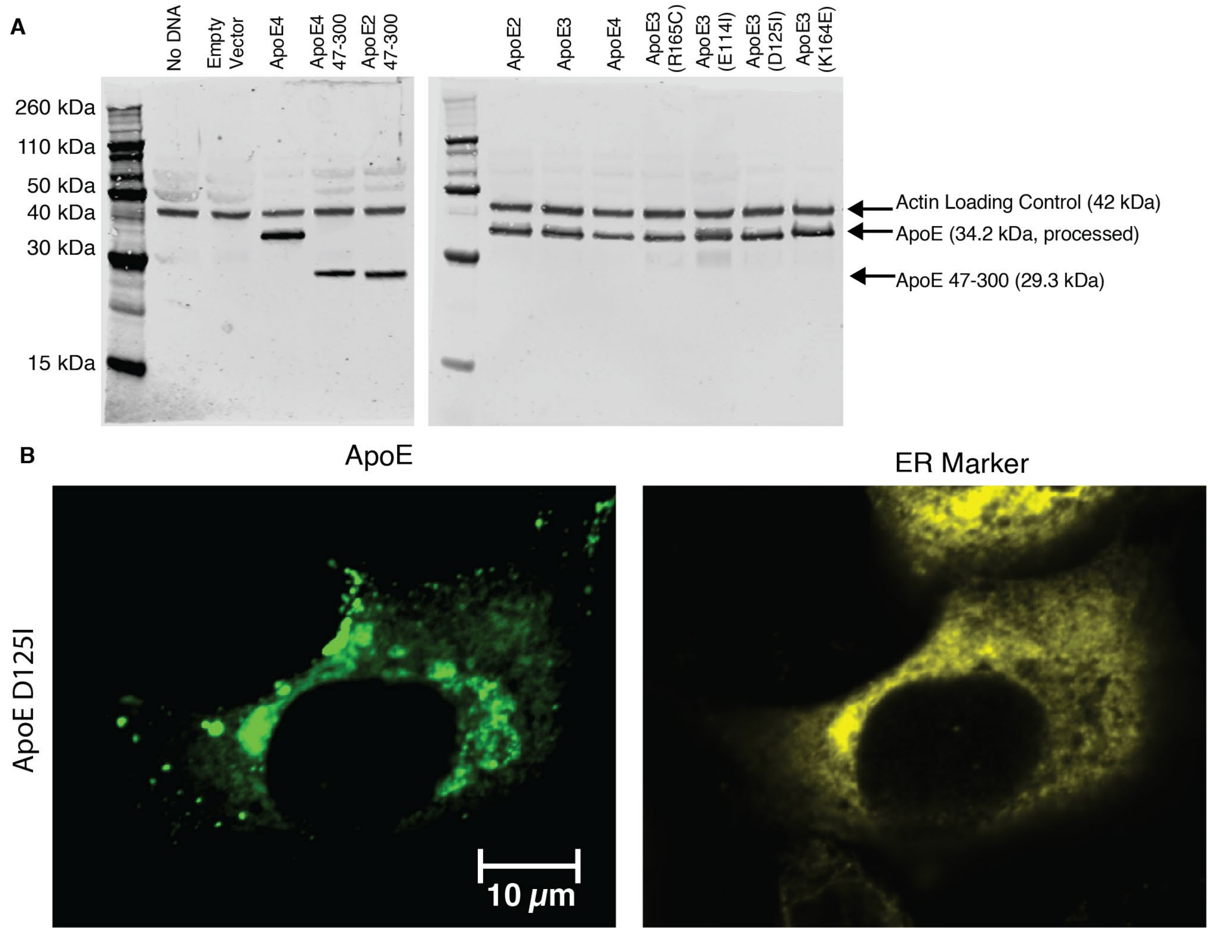


**Supplemental Figure 3** (A) Western blot data of some ApoE constructs showing expression at the expected molecular weight. (B) Sample images showing condensates forming in the full length ApoE3 D125I mutation.

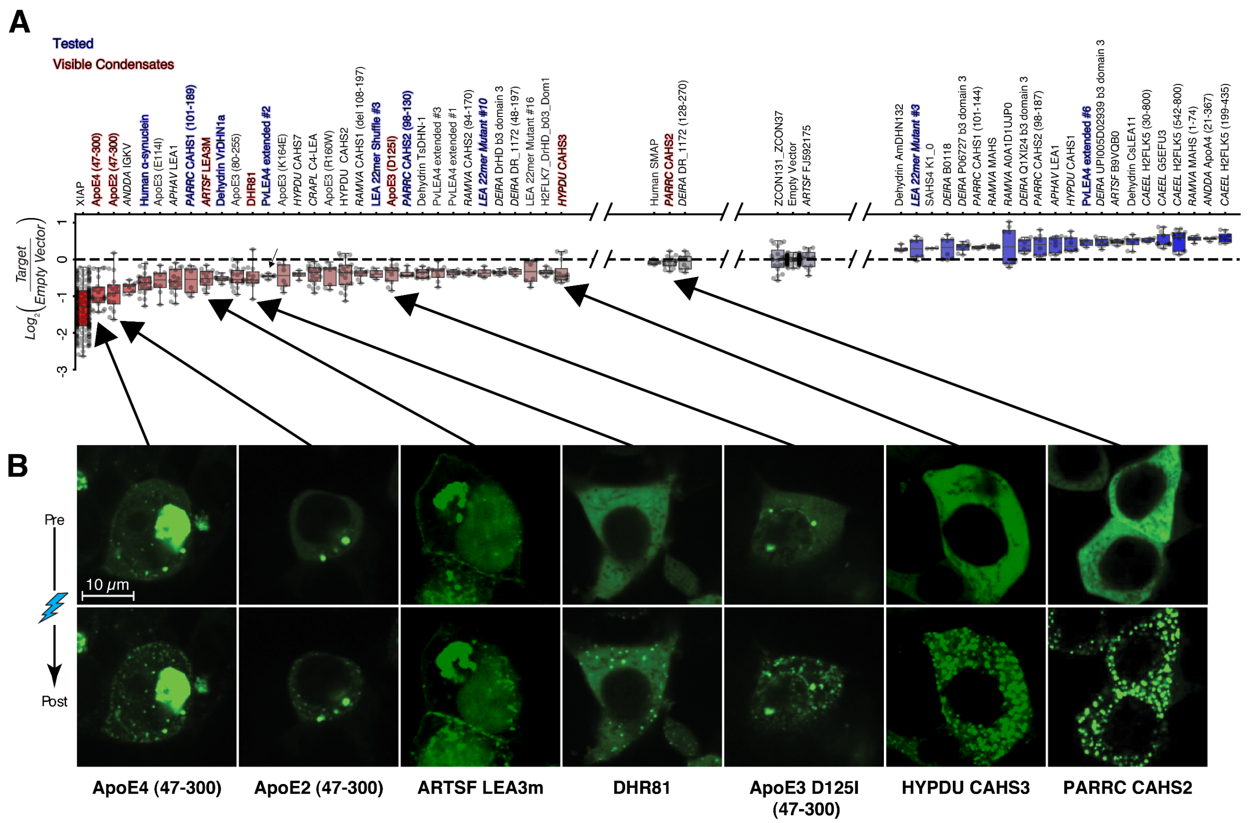


**Supplemental Figure 4** (A) Apoptosis assay data highlighting all the ExTol proteins tested and their apoptosis assay performance. Note that red and blue colored font refer to whether condensates were visible or not, respectively, via our microscopy. (B) Light-inducible condensate formation assays for selected ExTol proteins. Data shown includes all tested proteins with visible condensates.

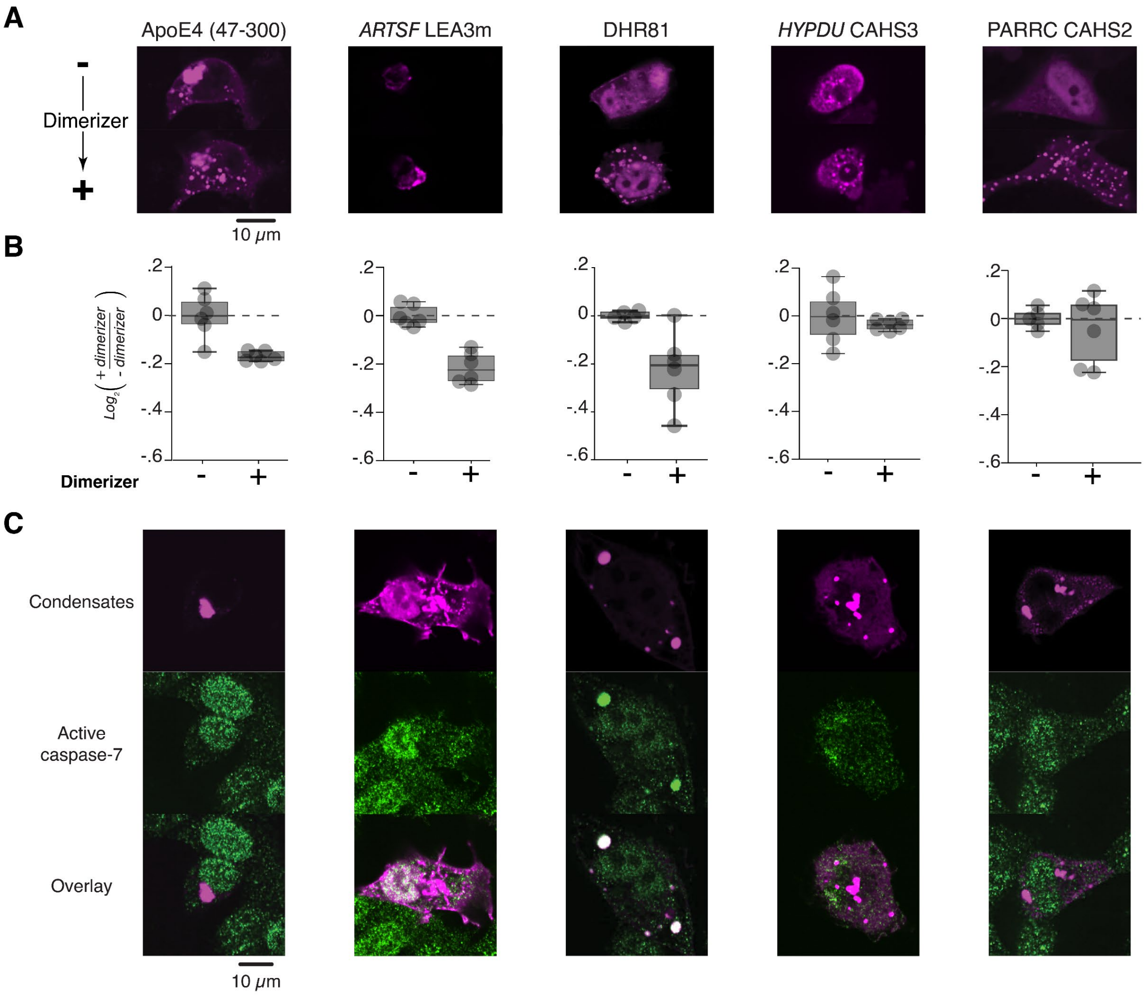
A
ApoE4 (47-300)
ARTSF LEA3m
DHR81
HYPDU CAHS3
PARRC CAHS2
-
Dimerizer
+
10 μm
B
Log2 (+ dimerizer / - dimerizer)
.2
0
-.2
-.4
-.6
Dimerizer
-
+
C
Condensates
Active caspase-7
Overlay
10 μm

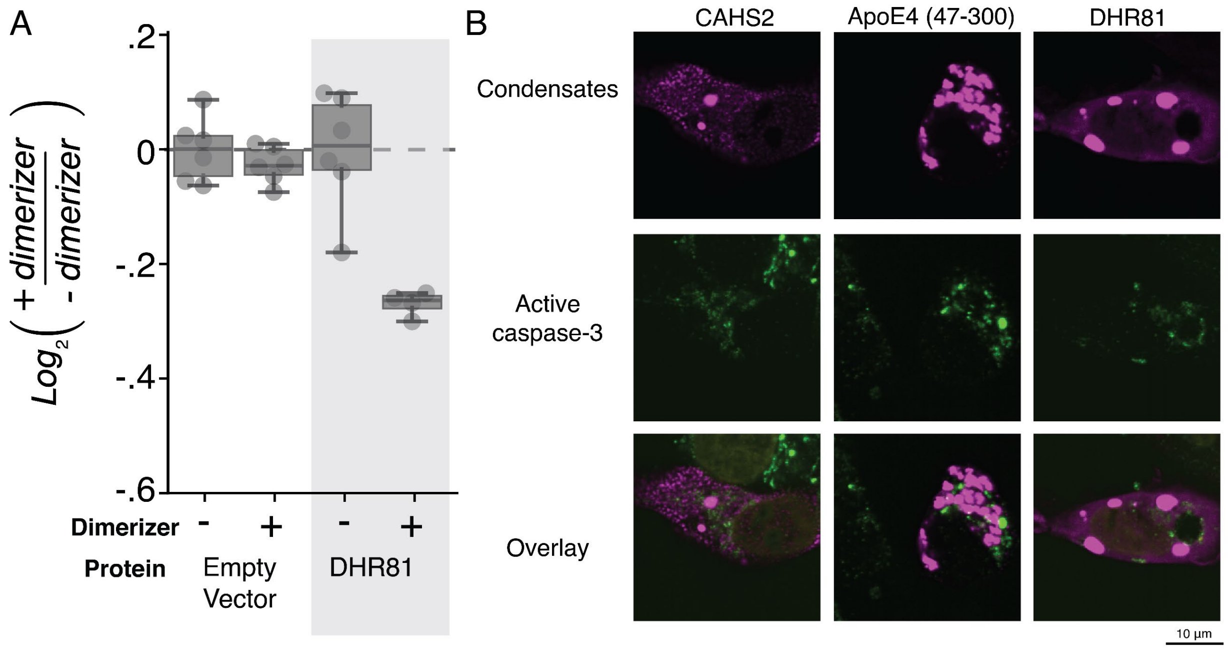


**Supplemental Figure 6** (A) DHR81 inducible condensates reduce apoptosis. (B) Sample images showing no co-localization between activated caspase-3 and condensates.

**Supplemental Table S1:** Supplemental table containing constructs and designs used in this study. See explanations tab for detailed descriptions of all the columns.

**Supplemental Table S2:** Data values obtained for apoptosis experiments. See explanations tab for detailed descriptions of all the columns.
