## Supplemental Methods for "Natural and designed proteins inspired by extremotolerant organisms can form condensates and attenuate apoptosis in human cells"

### *CONTACT FOR REAGENT AND RESOURCE SHARING*

Further information and requests for resources and reagents should be directed to and will be fulfilled by the Lead Contacts, Pamela Silver and Roger Chang.

### *EXPERIMENTAL MODEL AND SUBJECT DETAILS*

Escherichia coli strain DH5α (NEB) was used for all DNA synthesis applications and grown at 37°C in LB media with antibiotics. Mammalian cell culture work was done in Homo sapiens HT-1080. See methods below for more details.

### *METHOD DETAILS*

#### *Cell culturing and maintenance*

Eagle's Minimum Essential Medium (EMEM) (ATCC 20-2003), Heat-inactivated Fetal Bovine Serum (FBS) (Thermo Fisher Scientific 10082147), and Penicillin-Streptomycin (Corning 30-002-Cl) were all vacuum sterilized by filtration (0.22 µm pore size, Corning, 430767) and used for maintaining cell growth. Cells were maintained at 37°C. Homo sapiens HT-1080 cells (ATCC, CCL-121) were thawed in a 37°C water bath and immediately transferred to a cell culture flask. Cells were then grown for 48 hours. At this point, the media was gently aspirated and replaced and cells were allowed to grow for another 24 hours. Cells were then split into new flasks at one third of the density (Trypsin EDTA 1X 0.25% Trypsin/ 2.21mM EDTA in HBSS without sodium bicarbonate, calcium and magnesium, VWR 45000-664) and continued to grow in EMEM supplemented by FBS and Penicillin-Streptomycin.

#### *Cloning apoptosis constructs*

The empty vector used in the apoptosis assay was pSecTag2 A ordered from Thermo (V90020). A GFP sequence was cloned into this vector to provide a replacement sequence for Twist Bioscience to clone over. The resulting vector (YK 02, vector map can be found here https://benchling.com/s/seq-gjUPUvEDiLQT6K6CyGPl) was onboarded at Twist Bioscience. The cloning location was specified using the "replace GFP" feature in the YK 02 vector map described above. Column C of the "Designs" tab in Supplemental

Table S1 specifies the insert that was used to replace the GFP. The resulting vectors are identical to the YK 02 map shown above with the sequence from column C overwriting the “replace GFP” feature in the map.

*Cloning condensate constructs*

To generate the light-inducible constructs, the same YK 02 backbone was used. The target sequence was appended before a short linker, an mCherry sequence, another short linker, and the SspB protein (Addgene plasmid #122148). The chemically-inducible condensate sequences were generated by attaching the target protein up stream of an mCherry and 2x FKBP (F36V) sequences. The below table includes vector maps of the vectors created.

| Vector ID | Name | Sequence map |
|---|---|---|
| pBS1043 | CAHS2parrc-mCh-SspB | https://benchling.com/s/seq-mwhbZwTC9pJF1uR8owV2 |
| pBS1068 | ApoE2_47-300_mCh-SspB | https://benchling.com/s/seq-MCLamVEadIFGtsvDoeS0 |
| pBS1069 | ApoE4_47-300_mCh-SspB | https://benchling.com/s/seq-v0A5J1H806zWv64cogkG |
| pBS1072 | DHR81_Nccapped_mCh-SspB | https://benchling.com/s/seq-Yyqj5GsaPVjLGjLw7wUY |
| pBS1109 | CAHS2parrc-mCh-2xFKBP | https://benchling.com/s/seq-JahUmtrgF84VBOBCvEEj |
| pBS1111 | ApoE2_47-300_mCh-2xFKBP | https://benchling.com/s/seq-Nh0mlTXD5ciGEgfp7qCz |
| pBS1112 | ApoE4_47-300_mCh-2xFKBP | https://benchling.com/s/seq-geutb277CV3mCzZfKgOv |
| pBS1113 | ApoE3_47-300_mCh-2xFKBP | https://benchling.com/s/seq-vV2W6H7pDo4Itr6c85vW |
| pBS1114 | ApoE3_D125I_47-300_mCh-2xFKBP | https://benchling.com/s/seq-2ej7AZbyfB2FaIneJxdp |
| pBS1115 | DHR81_Nccapped_mCh-2xFKBP | https://benchling.com/s/seq-JFzThIQ8s83yDljKC0O6 |

*DNA Preparation*

All apoptosis constructs were synthesized by Twist Bioscience and transformed into DH5α cells for amplification and plasmid purification using the QIAGEN Plasmid Plus Midi Kit for transfection grade DNA purification.

*Cell seeding*

HT-1080 cells were washed with Phosphate-Buffered Saline (PBS) (Thermo Fisher Scientific, AM9625) and trypsinized (Trypsin EDTA 1X 0.25% Trypsin/ 2.21mM EDTA in HBSS without sodium bicarbonate, calcium and magnesium, VWR 45000-664). Cell density was determined by optical density with an automated cell counter (BIO RAD TC20), cell counting slides (BIO RAD, 1450015) and 0.4% solution Trypan Blue (BIO RAD, 1450021). After live cell count was obtained, cells were diluted to $7.5\times10^4$ cells per mL in Eagle's Minimum Essential Medium (EMEM) (ATCC, 20-2003) supplemented with 10% Heat-inactivated Fetal Bovine Serum (FBS) (Thermo Fisher Scientific, 10082147), that had been previously vacuum sterilized by filtration (0.22 µm pore size, Corning). 200 µL of cells were added to each well of a microscope grade 96 well plate (Corning 3603). Plates were then centrifuged at 330 rpm for 3 min (Eppendorf, Centrifuge 5810). Plates were then incubated at 37°C for 24 hours.

*Transfection*

HT-1080 cells were transfected 24 hours after plating on 96-well plates and were carried out in EMEM (ATCC, 20-2003) supplemented with 10% FBS (Thermo Fisher Scientific, 10082147). Prior to starting the transfection, the seeding media was removed and fresh media was added to the plate. Per well, 75 ng of target DNA and 25 ng of an mCherry transfection marker were transfected. This was achieved by mixing 360 ng of target DNA and 120 ng of mCherry transfection marker DNA together in a strip tube. Separately, a mix was prepared containing 750 µL of Opti-MEM (Thermo 31985062) and 30 µL of the p3000 reagent from the lipofectamine 3000 kit (Thermo Fisher Scientific, L3000015). 24 µL of this mix was added to the tube containing the target DNA and transfection marker. Again, in a separate tube, a mix was prepared containing 750 µL of Opti-MEM and 45 µL of the lipofectamine reagent. 24 µL of this mix was also added to the tube containing the target DNA and transfection marker. This complete mixture was briefly vortexed and allowed to stand for 5 minutes before 10 µL of the mix was added to each of the 4 wells on the plate.

*CPT treatment for caspase assay.*

24 hours after transfection, the transfection media was removed and replaced with FluoroBrite DMEM Media (Thermo A1896702) supplemented with 1x GlutaMAX (Thermo 35-050-061) and 10% FBS (Thermo Fisher Scientific, 10082147). This media also

contained 13.2 µM (S)-(+)-Camptothecin (Sigma C9911-100MG), 0.3 µM SiR-DNA stain (SpiroChrome CY-SC007), and 1.25 µM of the DEVD peptide for detecting caspase activation (Sartorius 4440). The plates were sealed with sterile Breathe-Easy film (USA scientific 9123-6100) and incubated at 37°C with 5% $CO_2$ for 30 minutes.

*Imaging for caspase assay without condensate induction*

After 30 min incubation, the plates were placed in an ImageXpress-confocal plate imager (Molecular Devices) at 37°C with 5% $CO_2$ and saturated humidity through the presence of a heated water bath. Images of all wells were taken every 45 minutes for 22.5 hours and stored on a remote server for later analysis.

*Imaging for caspase assay with condensate induction*

The imaging process was very similar to the caspase assay without condensate induction with the exception that most assays (those on plates 20210715, 20210914, and 20210924) were collected for 30 hours instead of 22.5. The images captured on the 20211226 plate were only captured for 22.5 hours.

*Western blotting*

Cells were cultured and transfected in 6 well dishes as described above but scaled up 30-fold to match the difference in surface area. The day after transfection, cells were scraped and collected into a 1.5 mL conical tube. Pellets were resuspended in about 1x pellet volume of RIPA buffer (10 mM Tris-Cl pH 8.0, 1 mM EDTA, 0.5 mM EGTA, 1% Triton X-100, 0.1% sodium deoxycholate, 0.1% SDS, 140 mM NaCl). Cells were allowed to lyse on ice for 30 min before being spun at 20,000xg for 1 min at 4°C. The resulting clarified lysate was run on 12% bis tris in MES buffer at 200 v for about 1 h. The gel was transferred to a membrane using the iBlot-2 membrane transfer system. The blot was then blocked for 1 hour at room temperature with agitation in 5% non-fat dried milk in TBS before adding the primary antibody and incubating overnight at 4°C. Below is a table of primary antibodies used and their dilution factors. All antibodies were used in TBST containing 1% non-fat dried milk.

| *Supplier* | *Order Number* | *Target* | *Dilution Factor* |
|---|---|---|---|
| Sigma | F1804 | FLAG | 1:100 |
| Abcam | ab24139 | ApoE | 1:200 |

| | | | |
|---|---|---|---|
| Abcam | ab8227 | Actin | 1:1000 |
| Abcam | ab8224 | Actin | 1:1000 |

The next day, the blot was washed 3x in TBST before being stained with secondary antibody for 1 hour at room temperature. Below is a table of the secondary antibodies used.

| *Supplier* | *Order Number* | *Target* | *Dilution Factor* | *Fluorophore* |
|---|---|---|---|---|
| VWR | 102971-154 | Mouse | 1:15000 | 800 CW |
| VWR | 103011-498 | Rabbit | 1:15000 | 680 RD |

Membranes were then scanned using a LICOR CLx instrument using auto exposure.

*Immunofluorescence*

Cells were grown on 96 well glass bottom plates from Cellvis (P96-1.5H-N). Cells were cultured and transfected as described above. Cells were fixed in 4% formaldehyde in culture medium. The cells were then washed 3x with PBS. Cells were then permeabilized with 100% cold MeOH for 10 min while being stored at -20°C. Permeabilized cells were blocked with PBS containing 1% BSA and 0.15% Triton X-100 for 1 hour. Blocking solution was removed and fresh blocking solution with a dilution of the primary antibody was added back to the well. Below is a table of the antibodies used and their dilution factors. Antibodies were left to bind overnight in the cold room.

| *Supplier* | *Order Number* | *Target* | *Dilution Factor* |
|---|---|---|---|
| Sigma | F1804 | FLAG | 1:500 |
| Thermo | MA3-019 | PDI | 1:400 |
| Abcam | ab24139 | ApoE | 1:500 |
| Cell Signaling Technologies | 9507S | Caspase-9 | 1:200 |
| Cell Signaling Technologies | 8438T | Caspase-7 | 1:500 |
| Cell Signaling Technologies | 9664T | Caspase-3 | 1:500 |
| Cell Signaling Technologies | 12963S | Cytochrome C | 1:200 |
| Cell Signaling Technologies | 89477S | Bax | 1:1000 |

| | | | |
|---|---|---|---|
| Cell Signaling Technologies | 2933S | Bim | 1:100 |
| Abcam | ab32445 | Bad | 1:50 |

The next day, the primary containing solution was removed, and the cells were washed with PBS 3x. PBS containing 1% BSA, 0.15% Triton X-100, and 2 µg/mL of DAPI was prepared. Depending on the source of the primary antibody and the desired fluorescent channel, the appropriate secondary antibody was added to stain the cells. This mix was added to the cells and allowed to stain for 1 hour in the dark at room temperature. Below is a table of the secondary antibodies used here.

| *Supplier* | *Order Number* | *Target* | *Dilution Factor* | *Fluorophore* |
|---|---|---|---|---|
| Thermo | A-11029 | Mouse | 1:2000 | Alexa Fluor 488 |
| Thermo | A-11008 | Rabbit | 1:2000 | Alexa Fluor 488 |
| Thermo | A-11031 | Mouse | 1:2000 | Alexa Fluor 568 |
| Thermo | A-11036 | Rabbit | 1:2000 | Alexa Fluor 568 |
| Thermo | A-21235 | Mouse | 1:2000 | Alexa Fluor 647 |
| Thermo | A-21245 | Rabbit | 1:2000 | Alexa Fluor 647 |

After staining, the cells were washed 3x with PBS before being stored in PBS for imaging.

*Confocal microscopy*

Confocal microscopy was done at the Nikon Imaging Center at Harvard Medical School using a Nikon Ti fluorescence microscope attached to a Yokogawa W1 spinning disk confocal. The microscope was equipped with a Nikon Perfect Focus System for focus assistance. A 60x objective was used for image capture (60x Nikon Plan Apo 60x 1.4 NA DCI N2). Live cell imaging was done with an Okolab Stage Top Incubator at 37°C and 5% $CO_2$ mounted on top of a Prior Proscan III motorized stage and shutters. Fixed cells were imaged in the PBS whereas live cells were imaged in FluoroBrite supplemented with 10% FBS and 1x GlutaMAX. The fluorophores used include the Alexa Fluor series (488, 568, and 647) as well as mCherry, DAPI, SiR-Hoechst (SpiroChrome, CHF260.00), and eGFP, depending on the experiment. Below is a table of the filters used to detect each fluorophore.

| Fluorophore | Excitation Laser | Emission Filter |
|---|---|---|
| Alexa Fluor 488 | 488 | Chroma ET525/50m |
| Alexa Fluor 568 | 561 | Chroma ET605/52m |
| Alexa Fluor 647 | 640 | Chroma ET705/72m |
| DAPI | 405 | Chroma ET455/50m |
| SiR-Hoechst | 640 | Chroma ET705/72m |
| mCherry | 561 | Chroma ET705/72m |
| eGFP | 488 | Chroma ET525/50m |

Excitation was done with a Nikon LUN-F XL solid state laser. Images were captured on a Hamamatsu ORCA-Fusion BT CMOS camera (6.5 $\mu m^2$ photodiode). Captured images were saved using NIS-Elements.

### *QUANTIFICATION AND STATISTICAL ANALYSIS*

#### *Image analysis for caspase assay*

The proprietary MetaXpress software was used to analyze the images. Within this software is a “find round objects” option. This was used to identify the initial nuclei of the cells. These nuclei were modified with the “expand without touching” option set to default settings. The area already defined by the nucleus was taken out using the area algebra feature to define the local area that is not the nucleus of the cell. The software was then set to quantify the FITC (DEVD peptide) and TRITC (Transfection marker) signals within the defined nuclear area and the local background. Each cell was assigned nuclear and background signals for TRITC and FITC values. These values were used to separate transfected from non-transfected cells and to quantify caspase activation.

#### *Quantification of apoptosis signal in transfected cells.*

Based on the image analysis described above, each cell was assigned a background subtracted transfection (TRITC) and apoptosis (FITC) value. This value was calculated by subtracting the local background signal from the nuclear signal to get the background subtracted signal. If the background subtracted transfection signal was above 300, then the cell was considered “transfected.” Otherwise, the cell was not considered for this analysis. The median apoptosis signal intensity was calculated by taking the median background subtracted FITC signal of all transfected cells in a particular well. This signal will heretofore be referred to as the apoptosis signal for the well.

*Data quality control.*

To ensure high quality data, an unequal variances two tailed t-test was performed comparing the XIAP positive control to the empty vector on each plate. If the t-test failed to reject the null hypothesis that the 4 XIAP wells had a lower apoptosis signal than the 4 empty vector wells, then the plate as a whole was discarded.

*Data Normalization and processing*

Each plate contained 4 replicates of each construct. The apoptosis signal for each well was calculated as described above. The 25$^{th}$ and 75$^{th}$ percentile was calculated. If the top or bottom points were more than 2 interquartile ranges away from these points, that point was identified as an outlier and removed. This value was then divided by an equivalent value from the empty vector run on the same plate (averaged across the 4 wells). The $Log_2$ of this value was calculated and will heretofore be referred to as the $Log_2$ Fold Change. Below is an equation showing how this value was calculated:

$$Log_2\ Fold\ Change\ =\ Log_2\left(\frac{Target\ Apoptosis\ signal}{Empty\ Vector\ Apoptosis\ Signal}\right)$$

Based on this analysis, the $Log_2$ Fold Change was calculated for each construct tested on each plate. This analysis amounts to 3 to 4 datapoints per date a construct was run.

*Data aggregation across days*

As each plate can only test a maximum of 21 constructs per plate with 4 wells per construct, data across dates were aggregated and analyzed for outliers. For a given construct, if a well had a $Log_2$ Fold Change more than 2 interquartile ranges away from the 25$^{th}$ or 75$^{th}$ percentile, then that well was removed. The resulting set of $Log_2$ Fold Changes for each construct are reported in “Construct level data” tab in Supplemental Table S2.

*Hydrophobic moment calculation*

Hydrophobic moment was calculated by assigning residue hydrophobicity based on the Fauchere-Pliska scale and then computing hydrophobic moment according to the formula defined in Eisenberg, 1982 [1-2]. Calculations assume a side-chain angular separation of 100°, typical of alpha helices. Python code to compute the hydrophobic moment was adapted from this script:

https://gist.github.com/JoaoRodrigues/568c845915aea3efa3578babfd72423c.

*PFAM domain analysis*

Protein sequences were analyzed for domain content using hmmscan (hmmer.org) to search sequences for PFAM-A domains [3]. Domain hits with full-sequence E-values < 0.01 were included in Supplemental Table S1 on the "pfam" analysis tab.

*HHRepID analysis*

Protein sequences were analyzed for repeat content using HHRepID [4]. An alignment was built for each protein using 3 iterations of HHblits against uniclust30 [5]. Then HHrepID was used with default parameters to determine repeats coverage. This analysis was used to highlight repetitive sequences that lack pfam domains in Figure 1.

*Disorder analysis*

Protein sequences were analyzed for disorder content using MobiDB-lite with default parameters [6].

### *Bibliography & References*


1. Fauchere, J.; Pliska, V., Hydrophobic parameters II of amino acid side-chains from the partitioning of N-acetyl-amino acid amides. *Eur. J. Med. Chem.* **1983,** *18*.
2. Eisenberg, D.; Weiss, R. M.; Terwilliger, T. C., The helical hydrophobic moment: a measure of the amphiphilicity of a helix. *Nature* **1982,** *299* (5881), 371-4.
3. Mistry, J.; Chuguransky, S.; Williams, L.; Qureshi, M.; Salazar, G. A.; Sonnhammer, E. L. L.; Tosatto, S. C. E.; Paladin, L.; Raj, S.; Richardson, L. J.; Finn, R. D.; Bateman, A., Pfam: The protein families database in 2021. *Nucleic Acids Res* **2021,** *49* (D1), D412-D419.
4. Biegert, A.; Soding, J., De novo identification of highly diverged protein repeats by probabilistic consistency. *Bioinformatics* **2008,** *24* (6), 807-14.
5. Steinegger, M.; Meier, M.; Mirdita, M.; Vohringer, H.; Haunsberger, S. J.; Soding, J., HH-suite3 for fast remote homology detection and deep protein annotation. *BMC Bioinformatics* **2019,** *20* (1), 473.
6. Piovesan, D.; Necci, M.; Escobedo, N.; Monzon, A. M.; Hatos, A.; Micetic, I.; Quaglia, F.; Paladin, L.; Ramasamy, P.; Dosztanyi, Z.; Vranken, W. F.; Davey, N. E.; Parisi, G.; Fuxreiter, M.; Tosatto, S. C. E., MobiDB: intrinsically disordered proteins in 2021. *Nucleic Acids Res* **2021,** *49* (D1), D361-D367.
